# Probing the Plasticity in the Active Site of Protein N-terminal Methyltransferase 1 Using Bisubstrate Analogs

**DOI:** 10.1101/2020.04.13.039990

**Authors:** Dongxing Chen, Cheng Dong, Guangping Dong, Karthik Srinivasan, Jinrong Min, Nicholas Noinaj, Rong Huang

## Abstract

The interactions of a series of bisubstrate analogs with protein N-terminal methyltransferase 1 (NTMT1) were examined to probe the molecular properties of the NTMT1 active site through biochemical characterization and structural studies. Our results indicate that a 2-C to 4-C atom linker enables its respective bisubstrate analog to occupy both substrate and cofactor binding sites of NTMT1, but the bisubstrate analog with a 5-C atom linker only interacts with the substrate binding site and functions as a substrate. Furthermore, the 4-C atom linker is the optimal and produces the most potent inhibitor (*K*_i, app_ = 130 ± 40 pM) for NTMT1 to date, displaying over 100,000-fold selectivity over other methyltransferases and 3,000-fold even to its homolog NTMT2. This study reveals the molecular basis for the plasticity of the NTMT1 active site. Additionally, our study outlines a general guidance on the development of bisubstrate inhibitors for any methyltransferases.

## Introduction

Protein methylation, a widespread epigenetic modification that is catalyzed by protein methyltransferases (PMTs), plays critical roles in many cellular processes such as gene expression, transcriptional regulation, DNA replication and damage repair. Dysregulation of PMT expression has been implicated in a variety of diseases including cancer, cardiovascular, neurodegenerative, and other diseases.^[1]–[5]^ Therefore, there have been emerging interests in development of selective inhibitors for PMTs as potential therapeutic agents. Recently, several inhibitors of PMTs have been developed and entered human clinical trials, such as DOT1-like protein inhibitor EPZ-5676 (Phase I/II), protein arginine methyltransferase 5 inhibitor GSK3326595 (Phase I/II), enhancer of zeste homolog 2 inhibitors EPZ-6438 (Phase I/II) and CPI-1205 (Phase I/II).^[2]^

PMTs utilize the cofactor S-adenosyl-L-methionine (SAM) as a methyl donor to transfer the methyl group from SAM onto the substrate with concomitant generation of S-adenosine-L-homocysteine (SAH) and methylated substrate.^[2]^ The PMT-catalyzed methylation reaction typically undergoes a Bi-Bi mechanism and involves the formation of a ternary complex.^[6]^ Bisubstrate analogs that occupy both substrate and cofactor binding sites have the potential to achieve higher binding affinities than a substrate analog.^[7]–[10]^. Furthermore, bisubstrate analogs may offer superior selectivity for their targeted enzyme.^[9],[11],[12]^ On account of a shared SAM cofactor binding pocket across PMT families, it is rational to hypothesize that bisubstrate analogs can be a general approach to offer potent and specific inhibitors for each individual PMT. Recent progresses have been made in the development of bisubstrate analogs for a few PMTs, such as protein N-terminal methyltransferase 1 (NTMT1)^[11],[13],[14]^, coactivator-associated arginine methyltransferase 1 (CARM1)^[15]^, protein arginine methyltransferase 1 (PRMT1)^[16]^, and PRMT6^[17]^. These bisubstrate analogs were created by connecting a SAM analog to a substrate or a functional group of the corresponding substrate. However, the potency of those bisubstrate analogs for their respective targets varied from nanomolar to micromolar. This serendipitous potency imposed the need of explicit instruction on design of bisubstrate analogs for PMTs.

The aforementioned need has prompted us to investigate how the linker length would affect the interaction of bisubstrate analog to its target PMT. Here, we used NTMT1 as a model system to examine the optimal length of the linker, with the intention to provide a general guide to facilitate the design of optimal bisubstrate analogs for other PMTs. NTMT1 recognizes a unique X-P-K/R motif at the protein alpha-N-terminus.^[18]–[20]^ It methylates centromere-specific histone H3 variants centromere protein A/B, regulator of chromatin condensation 1, damaged DNA-binding protein 2 and poly(ADP-ribose) polymerase 3.^[21]–[26]^ Hence, NTMT1 has been implicated in regulating chromosome segregation, mitosis, and DNA damage repair.^[22],[24],[27]^ We have previously developed bisubstrate analogs for NTMT1. Among them, compound NAH-C3-PPKRIA showed an IC_50_ value of 35 ± 2 nM in a MALDI-MS based assay, and displayed more than 100-fold selectivity against other methyltransferases (Figure 1A).^[11]^ In addition, compound NAH-C3-GPRRRS showed an IC_50_ value of 0.94 ± 0.16 *μ*M for NTMT1.^[14]^ Both compounds have a propylene linker, which raises the question if a propylene linker is optimal. To address this question, we design and synthesize a series of NTMT1 bisubstrate analogs with variable linker lengths to probe the active site of NTMT1.

**Figure 1.**
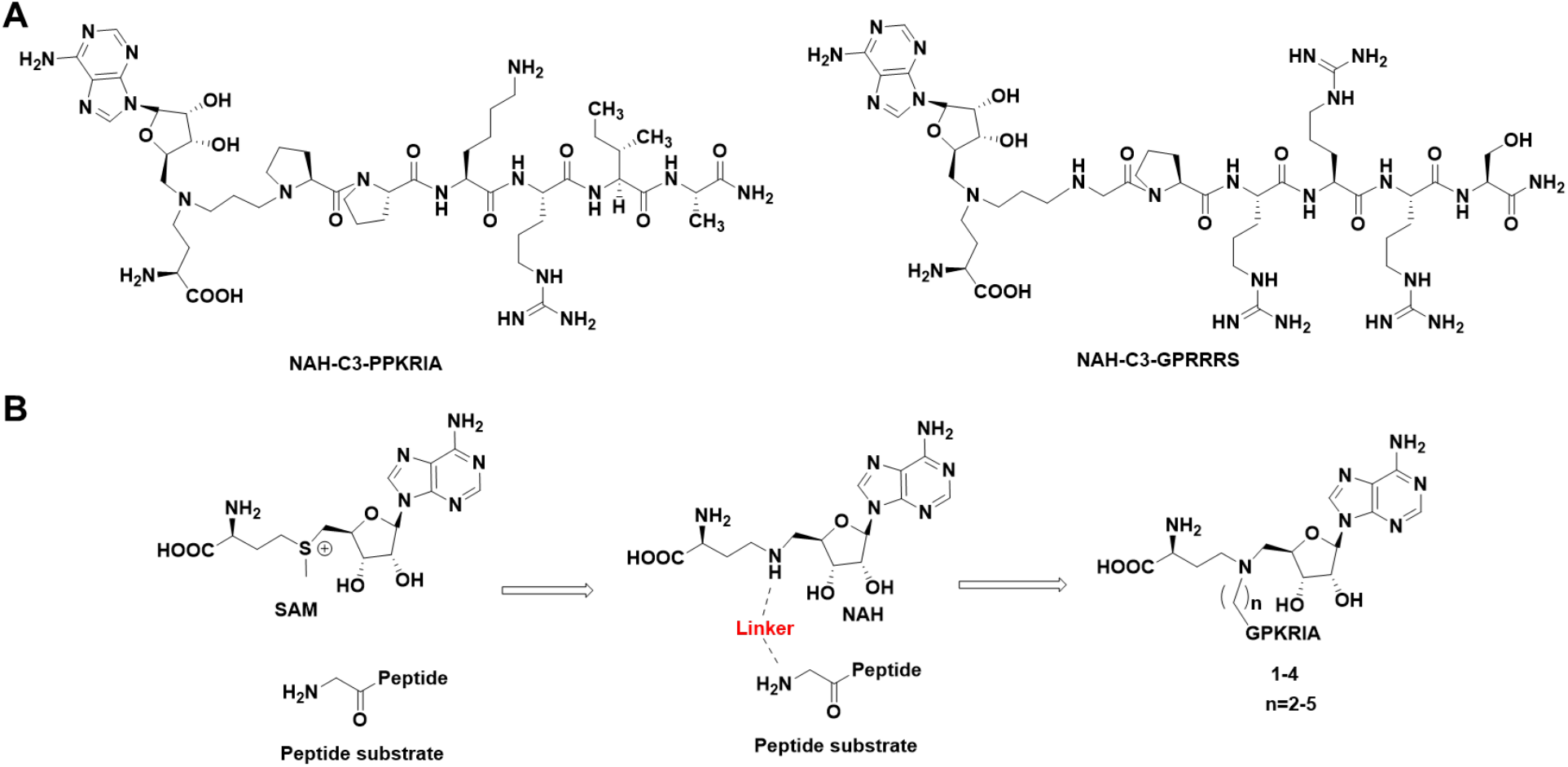
(A) Chemical structures of NTMT1 inhibitors. (B) Design of the bisubstrate analogs for NTMT1.

## Results

The linear distance between the sulfur atom of SAH and the nitrogen atom of substrate ranges 4.6 Å to 5.5 Å in the ternary structures of NTMT1.^[18],[19]^ Therefore, we varied the linker from 2 to 5 methylene groups (Figure 1B, Scheme 1). We chose the peptide GPKRIA due to the synthetic versatility of Gly at the first position. Compounds **1-4** were synthesized in a convergent manner by reacting the primary amines **8-11** with α-bromo peptide on resin and followed by acid treatment (Scheme 1).^[14]^

**Scheme 1.**
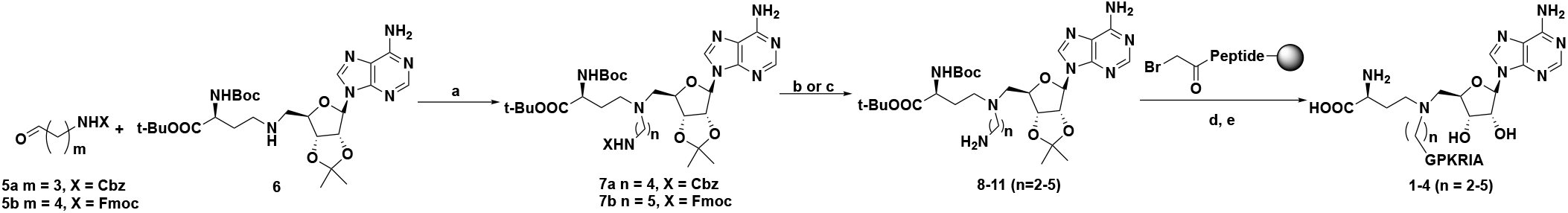
Synthetic route for compounds 1-4. Reagents and conditions: a) NaBH_3_CN, CH_3_OH, r.t., 38-68%; b) 10% Pd/C, H_2_, MeOH, 56%; c) DBU, CH_3_CN, 0 °C to r.t., 71%; d) α-bromo peptides, DIPEA, DMF, r.t.; e) TFA: DODT: H_2_O: TIPS = 94: 2.5: 2.5: 1; f) TFA, CH_2_Cl_2_, 0 °C to r.t., 42%; g) 4N HCl in dioxane, H_2_O, 0 °C to r.t., 45%.

**Figure 2.**
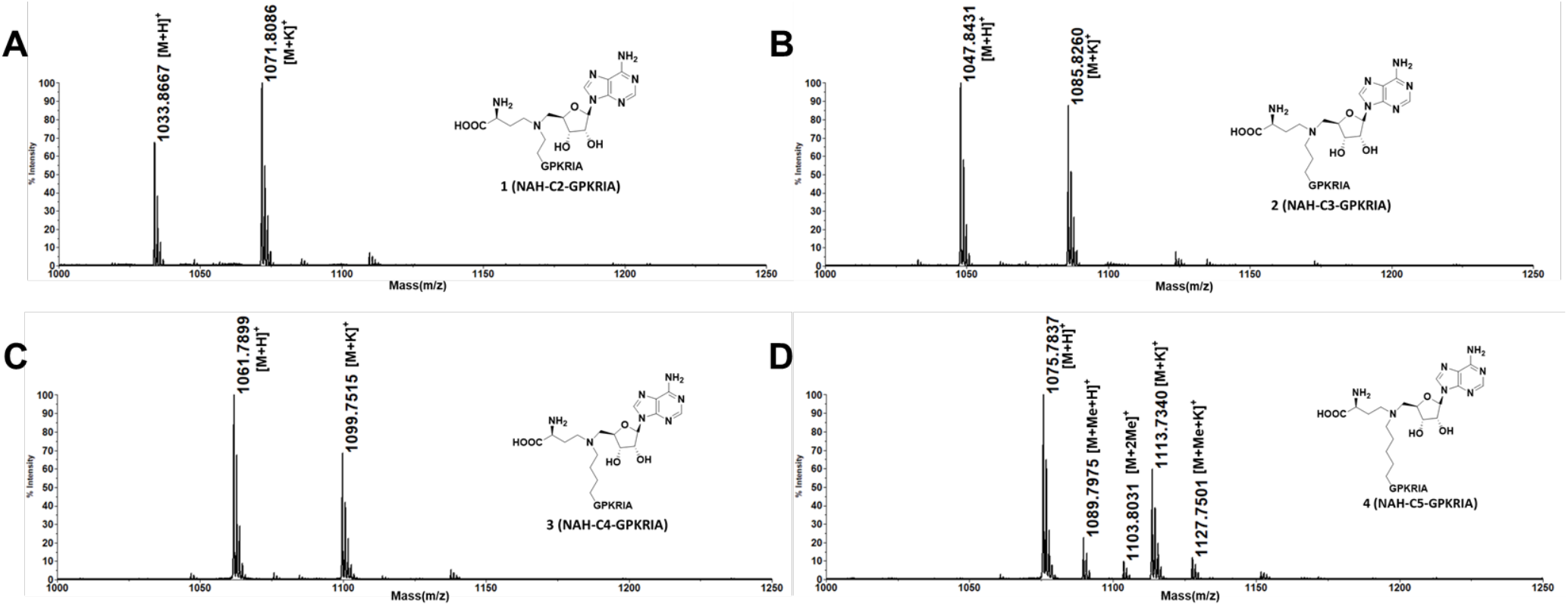
MALDI-MS methylation assay for the bisubstrate analogs **1-4**.

Compounds **1-4** were first subject to methylation assay catalyzed by NTMT1 in the presence of SAM to first check if they may act as possible substrates. Surprisingly, compound **4** containing the 5-C atom linker was able to be methylated by NTMT1 (Figure 2), indicating **4** as a substrate for NTMT1. Then, the inhibitory activities of **1-3** were evaluated in the SAH hydrolase (SAHH)-coupled fluorescence assay with 10 minutes of preincubation time.^[28]^ Both SAM and the peptide substrate GPKRIA were at their respective *K*_m_ values. In order to compare the inhibition activities of NAH-GPKRIA series under the same condition, we resynthesized and purified **1** and **2** with an ethylene and a propylene linker, respectively. Their IC_50_ values were lower than previous values^[14]^, which may be caused by an improved S/N ratio with a different GPKRIA peptide substrate and fluorescent dye ThioGlo4. The measured IC_50_ values for most compounds converged to the enzyme concentration (100 nM) present in the assay (Table S1), suggesting tight-binding inhibition. Therefore, using Morrison’s equation to define their apparent *K*_i_ values is a better method to analyze the concentration-response data for these tight binding inhibitors.^[29],[30]^. The tight-binding inhibitors often exhibit slow binding traits, but Morrison’s equation applies to the steady state velocity.^[29],[30]^ Therefore, compounds **1-3** were chosen to investigate the effects of preincubation time on the inhibitory activity of tight-binding inhibitors (Figure 3). The *K*_i,app_ values of the three compounds were inversely proportional to the pre-incubation time (Figure 3B, D and F), indicating their slow binding feature. The results showed the interaction between the compounds and NTMT1 almost reached equilibrium after 30 min of incubation. Compound **3** containing a 4-C atom linker displayed the lowest *K*_i,app_ value (130 ± 40 pM) (Figure 3B) after 120 min, further substantiating its slow and tight binding feature ([E]/*K*_i,app_ > 100).^[29],[30]^ When the linker length decreased from a 4-C atom to a 3-C and 2-C, compound **2** (*K*_i,app_ = 1.3 ± 0.6 nM) and **1** (*K*_i,app_ = 5.4 ± 2.6 nM) exhibited a 10-fold and 40-fold decrease in comparison with **3**, respectively.

NTMT2 is a closely-related homolog of NTMT1, which shares over 50% sequence similarity and the same X-P-K recognition motif.^[31]^ To assess the interactions of the bisubstrate anlogs with both enzymes, the binding affinities of **1-4** for both NTMT1 and NTMT2 were determined by isothermal titration calorimetry (ITC) experiment (Figure 4). All four compounds bound to NTMT1, especially compound **3** (K,_d1/d2_ = 0.6/0.3 *μ*M). Notably, the *K*_d_ values of bisubstrate analogs were much higher than their *K*_i, app_ values except compound **4**. This difference may be caused by endogenous amounts of SAM/SAH binding to recombinant NTMTs as reported before.^[18],[31]^ However, the ITC binding data was still informative, since the bisubstrate analogs at 1 mM showed negligible interactions with NTMT2.^[31]^

**Figure 3.**
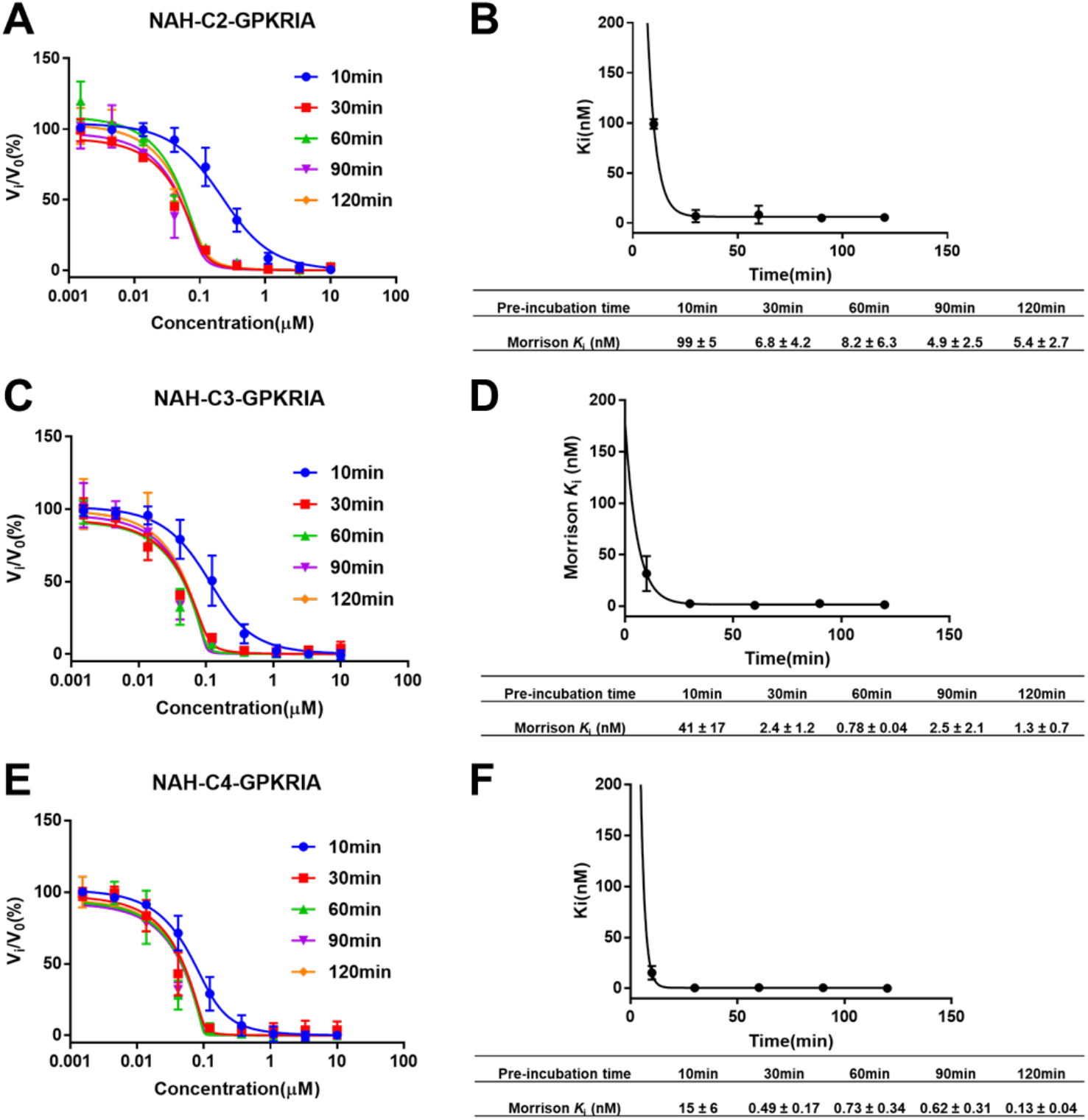
Time-dependent inhibition study of compounds **1-3**. All experiments were run with duplicate (n=2), except 10min of pre-incubation (n=3). (A), (C) and (E) Concentration-response plots for different pre-incubation times fitted to Morrison’s quadratic equation for compounds **1**, **2** and **3**, respectively; (B), (D) and (F) Nonlinear regression of Morrison *K*_i, app_ values with pre-incubation time for compounds **1**, **2** and **3**, respectively.

To further confirm the inhibitory activity of bisubstrate inhibitor, the MALDI-MS methylation assay was performed as an orthogonal assay to monitor the production of methylated peptides with both substrate peptide GPKRIA and SAM at their *K*_m_ values after 10 min of preincubation time (Figure 5A-B). At 50 nM of compound **3**, tri-methylation of GPKRIA peptide was substantially reduced. At 200 nM of compound **3**, tri-methylation of GPKRIA peptide was barely detected (Figure 5B).

**Figure 4.**
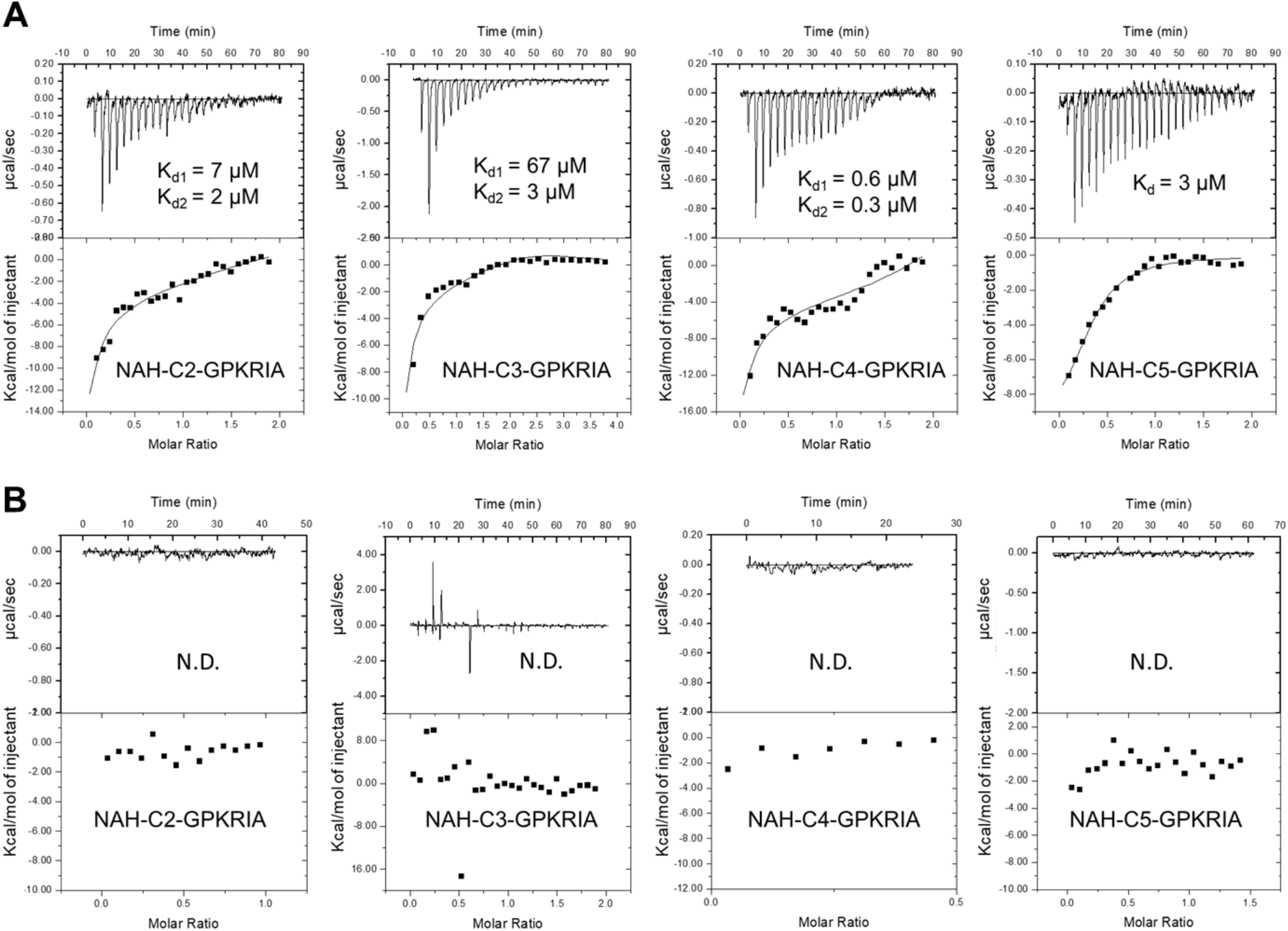
Binding affinity for NTMT1(A) and NTMT2 (B). N.D.: not detected.

**Figure 5.**
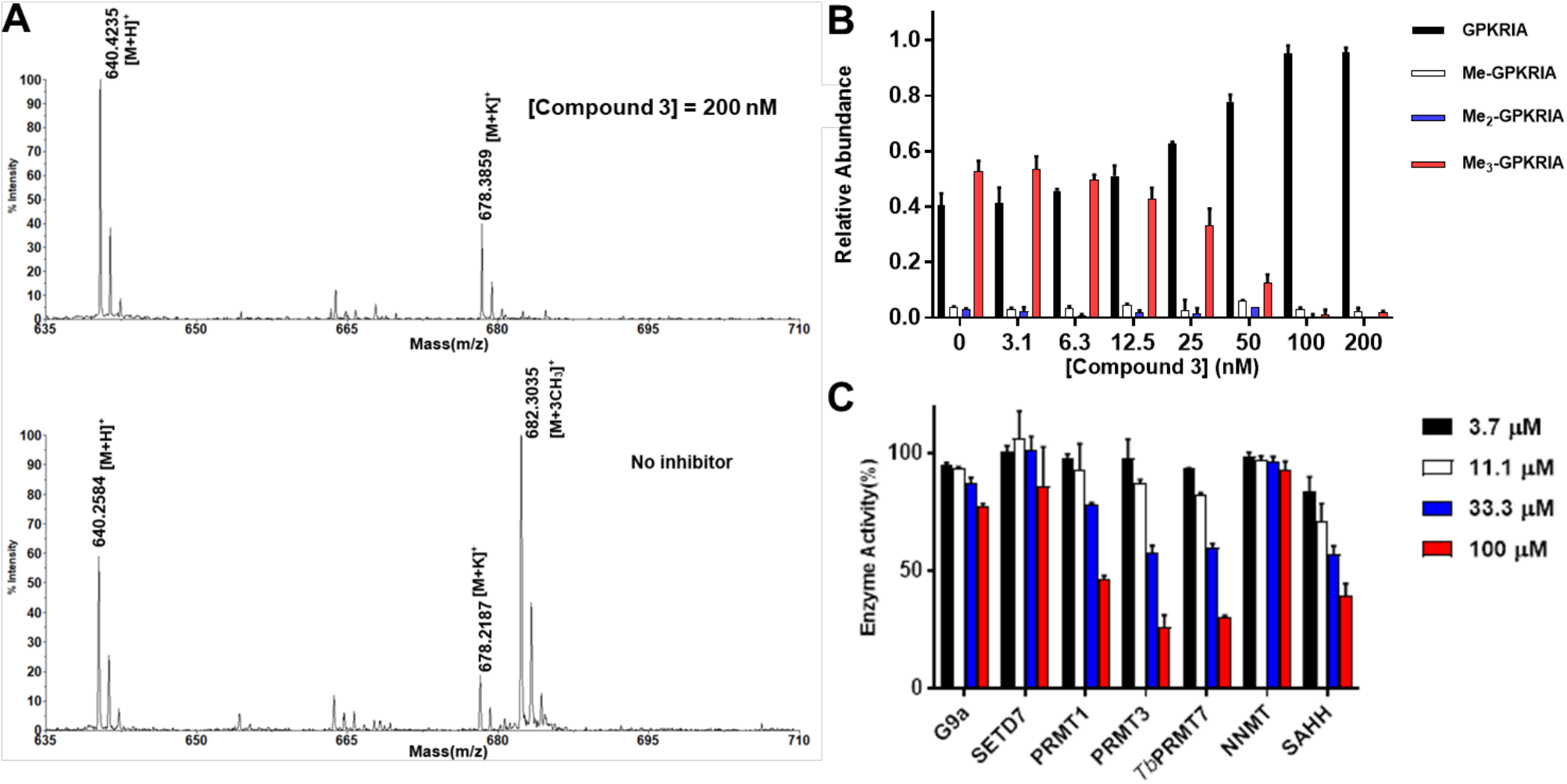
(A) and (B) MALDI-MS methylation inhibition assay compound **3** (n=2). (A) MALDI-MS results of MALDI-MS methylation inhibition assay for compound **3**; (B) Quantification of methylation progression of GPKRIA by NTMT1 with **3** at 20 min. (C) Selectivity evaluation of compound **3** (n=2).

The selectivity of **3** for NTMT1 was also examined against a panel of methyltransferases that shared a SAM binding pocket, which also included the coupling enzyme SAHH that has a SAH binding site and was used in the fluorescence assay (Figure 5C, Table S2). The compound barely showed inhibitory activity over euchromatic histone-lysine N-methyltransferase 2 (G9a), SET domain-containing protein 7 (SETD7) and nicotinamide N-methyltransferase (NNMT) at 100 *μ*M. At 33 *μ*M, **3** demonstrated less than 50% inhibition over protein arginine methyltransferase 1 (PRMT1), protein arginine methyltransferase 3 (PRMT3), *Trypanosoma brucei* protein arginine methyltransferase 7 (*Tb*PRMT7) and SAHH. Therefore, compound **3** showed more than 100,000-fold selectivity over other methyltransferases.

To understand the molecular basis of the effect of different linkers, we obtained the co-crystal structures of bisubstrate analogs **1** and **4** with NTMT1 (Figure 6). The binary complex X-ray crystal structure of NTMT1-**1** complex (PDB: 6WJ7) illuminates the inhibition mechanism since compound **1** does occupy both cofactor and substrate binding sites of NTMT1. It also overlays well with previously reported bisubstrate inhibitor **NAH-C3-PPKRIA** in the X-ray crystal structure of NTMT1-**NAH-C3-PPKRIA** (PDB code: 6DTN) (Figure 6B). The SAM analog moiety of **1** in the binary complex binds almost same as **NAH-C3-PPKRIA**, which is nearly identical with SAH in the ternary complex of substrate peptide/SAH with NTMT1. When increasing the carbon linker to 5-C atom linker, the peptide part of the compound **4** retained its interaction with the substrate binding pocket and overlays very well with the PPKRIA peptide portion of **NAH-C3-PPKRIA** (Figure 6D-E). However, the NAH moiety of the compound **4** no longer occupied the SAM binding pocket. Interestingly, the SAM analog part inserted to a new pocket adjacent to the substrate binding pocket, which resulted in exposure of the secondary amine (α-N-terminal amine of the peptide portion) of compound **4** to receive the methyl group from SAM. Nevertheless, it infers the length of 5-C atom linker is too long to allow the bisubstrate analog **4** to simultaneously occupy both SAM and peptide binding sites.

**Figure 6.**
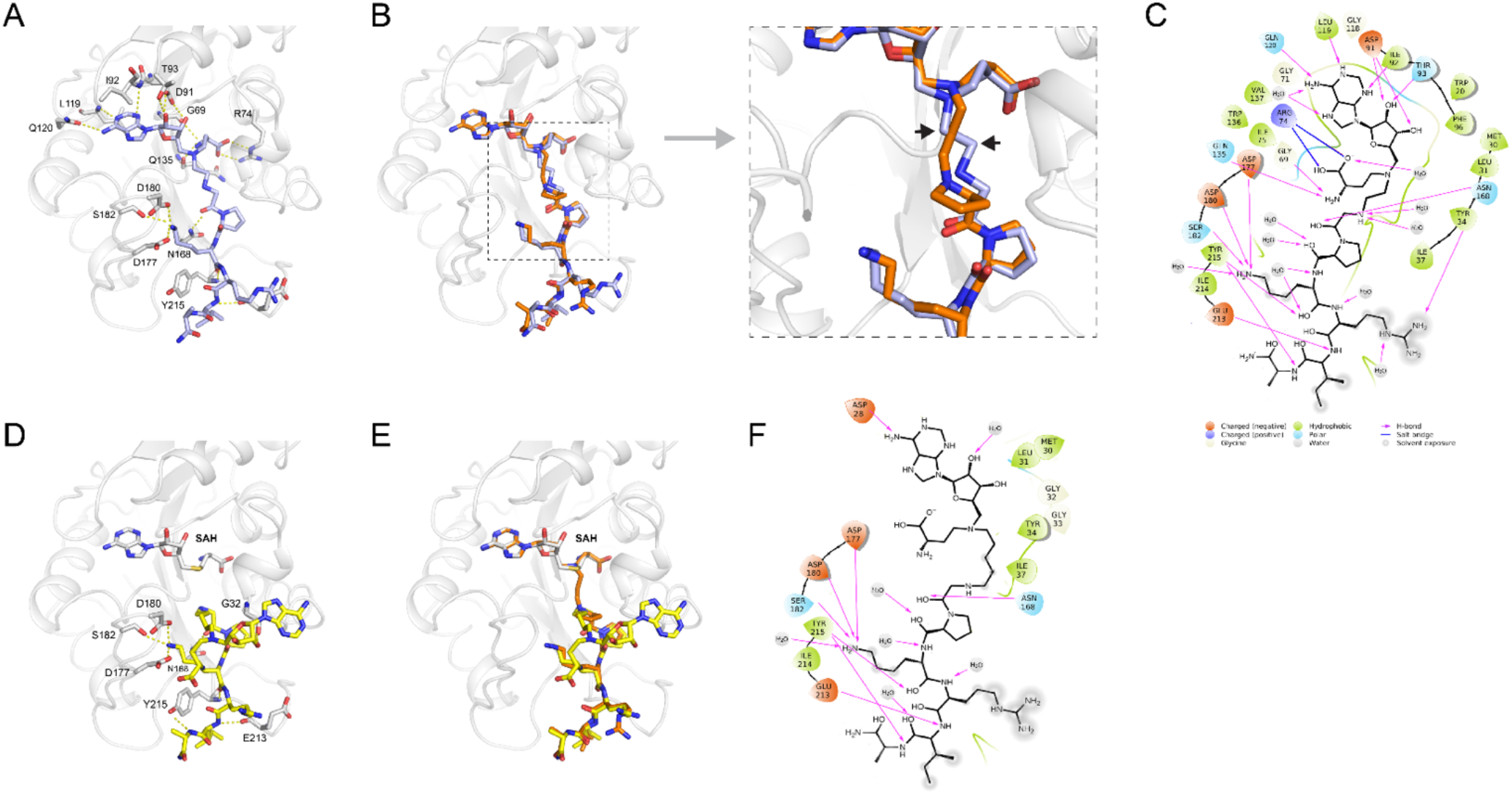
X-ray co-crystal structures. (A) The X-ray crystal structure of compound **1** (purple stick) in complex with NTMT1 (gray ribbon) (PDB code: 6WJ7). H-bond interactions are shown in yellow dotted lines; (B) Overlay of the crystal structure of NTMT1(gray ribbon)-**1** (purple stick) complex with the crystal structure of the NTMT1-NAH-C3-PPKRIA (orange stick) binary complex (PDB code: 6DTN); (C) Compound **1** 2D interaction diagram (Schrödinger Maestro) with NTMT1; (D) The X-ray crystal structure of NTMT1(gray ribbon)-**4** (yellow stick)-SAH (gray stick) (PDB code: 6PVB). H-bond interactions are shown in yellow dotted lines; (E) Overlay of the crystal structure of NTMT1(gray ribbon)-**4** (yellow stick) - SAH (gray stick) complex (PDB code: 6PVB) with the crystal structure of NTMT1-NAH-C3-PPKRIA (orange stick) binary complex (PDB code: 6DTN).

### Inhibition Mechanism Studies

Despite additional efforts to screen and optimize the conditions, cocrystal structure of NTMT1 in complex with **2** or **3** was not obtained. Hence, Inhibition mechanism studies were performed to confirm the binding modes for 4-C atom linker compound. Since the co-crystal structures of NAH-C3-PPKRIA containing the 3-C linker revealed that the compound could simultaneously occupy SAM and peptide binding sites, it was reasonable to assume that compound **2** with 3-C linker could also interact with both SAM and peptide binding sites. So, inhibition mechanism studies for compound **3** was performed along with compounds **1** and **2** as the control under the condition of 2h of pre-incubation (Figure 7). Even at both 6 *K*m and 12 *K*m concentrations of peptide substrate, there were slight competition for compound **3**. Despite barely competition for compounds **2** and **3**, ~10% of enzymatic activity was rescued in presence of 0.3 *μ*M or higher concentration of inhibitors. When competing peptide substrate concentration increased to 50 *K*m, slightly right shift was observed for **2** and **3**. In addition, ~20% enzymatic activity was rescued, further reflecting their tight and slow-binding character. When increasing the concentration of SAM from 6 *K*m to 25*K*m, similar trends were observed. The catalytic activity of NTMT1 could be rescued with higher concentration of SAM or peptide, suggesting the compounds could be competitive with peptide and SAM. Those results indicated that all three bisubstrate inhibitors **1-3** occupied both SAM and peptide substrate binding sites in a reversible manner.

**Figure 7.**
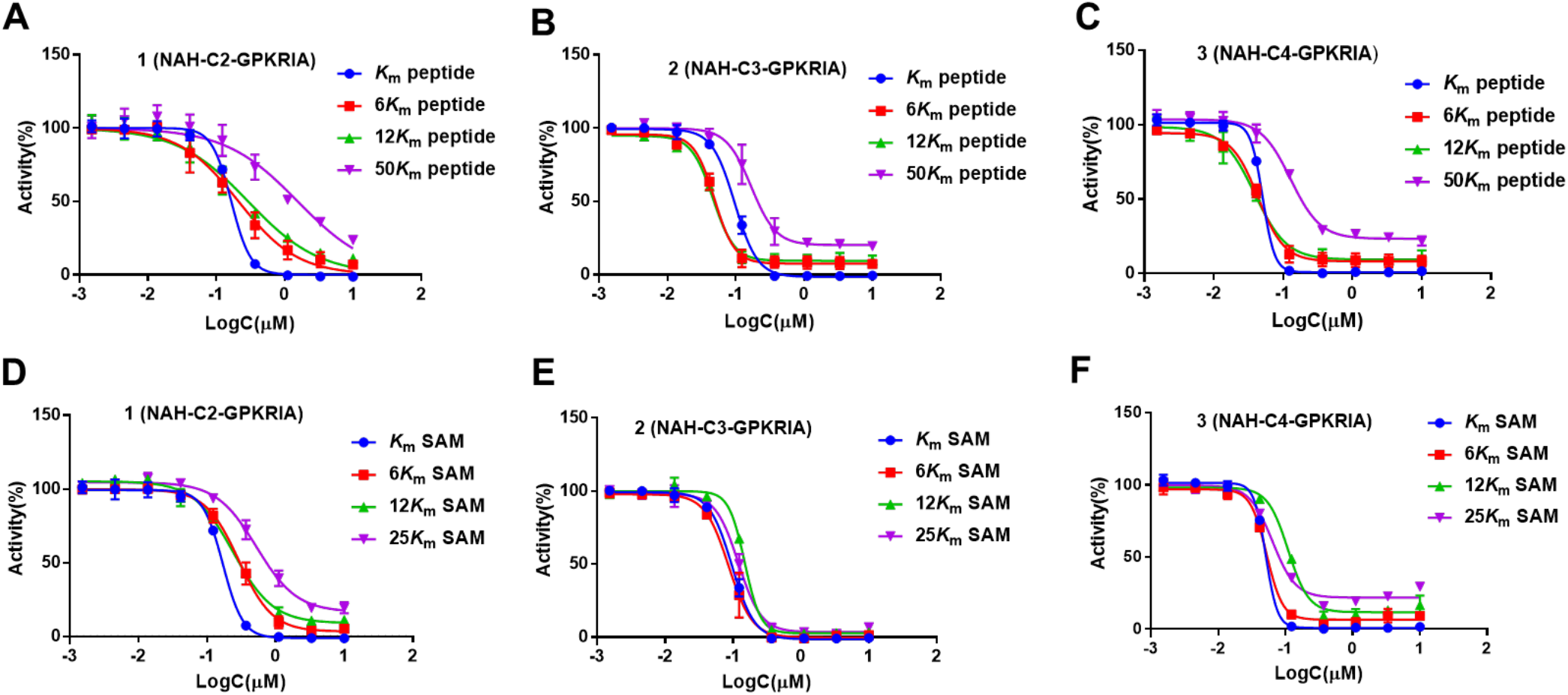
Inhibition Mechanism Studies for compounds **1-3**. (A-C) IC_50_ curves of **1-3** at varying concentrations of peptide with fixed concentration of SAM; (D-F) IC_50_ curves of **1-3** at varying concentrations of SAM with fixed concentration of peptide. The pre-incubation time for the assay was 120 min, and all the experiment were run in duplicates (n = 2).

## Discussion

This series of tight-binding and selectice bisubstrate analogs **1**-**4** to NTMT1 represent valuable tools to offer valuable insights into the astounding plasticity in the transition state of NTMT1 since it can accommodate linker length from 2-C to 4-C atom. Among them, a 4-C atom linker is the optimal length to achieve the best inhibition with a *K*_i, app_ value of 130 pM. Notably, a 5-C atom linker would convert a tight-binding bisubstrate analog to a substrate for NTMT1, suggesting the upper limit of the aforementioned plasticity. The co-crystal structure of **4** in complex with NTMT1 (PDB code: 6PVB) implied that extending the linker to a 5-C atom linker may cause tension to prevent the bisubstrate analog from engaging with both binding sites. Given the transient nature of transition state, these bisubstrate analogs provide dynamic snapshots of the reorganization within the NTMT1 active site during the catalysis.

Furthmore, top inhibitor **3** (NAH-C4-GPKRIA) containing a 4-C atom linker showed picomolar inhibition (*K*_i, app_ = 130 ± 40 pM) for NTMT1, and more than 100,000-fold selectivity over other MTases. In addition, all bisubstrate analogs **1**-**3** demonstrate no interaction with its close homolog NTMT2 despite of the similar catalytic sites of NTMT1/2.^[31]^ This delicate selectivity further corroborates the benefits of bisubstrate analogs on account of their high specificity to offer tight and selective inhibitors for individual target. However, increasing the carbon linker length to a 5-C atom resulted in a substrate instead of an inhibitor for NTMT1, which defined the scope or limit of plasticity. To our knowledge, this is the first case that delineates the optimal linker range of MTase bisubstrate analog and provide the molecular basis for the obliterated activity for longer linkers.

The linear distance between the sulfonium ion and the nitrogen atom that accepting methyl group is very similar for most PMTs, which ranges from 3.5 Å to 5.5 Å among the ternary complex of all examined PMTs including NTMT1, SET domain-containing protein 8, and euchromatic histone-lysine N-methyltransferase 1.^[18],[19],[32]–[35]^ Therefore, we propose that a 3-C or 4-C atom linker would serve as a general and feasible starting point to build bisubstrate analogs for any MTase. The optimal linker length has also been corroborated by successful inhibitors for RNA MTase, as a 3-C atom linker showed better inhibition compared to a 2-C atom linker.^[36]^ Future directions will focus on applying such linkers to conjugate small molecule inhibitors that target individual binding sites to achieve higher selectivity and cellular potency.

## Materials and Methods

### Chemistry General Procedures

The reagents and solvents were purchased from commercial sources (Fisher) and used directly. The intermediates and products were purified by using an ISCO CombiFlash system and prepacked SiO_2_ cartridges. Final compounds were purified on preparative high-pressure liquid chromatography (RP-HPLC) was performed on Agilent 1260 Series system. Systems were run with 0-20% methanol/water gradient with 0.1% TFA. NMR spectra were acquired on a Bruker AV500 instrument (500 MHz for ^1^H-NMR, 126 MHz for ^13^C-NMR). TLC-MS were acquired using Advion CMS-L MS. Matrix-assisted laser desorption ionization mass spectra (MALDI-MS) data were acquired in positive-ion mode using a Sciex 4800 MALDI TOF/TOF MS. The peptides were synthesized using a CEM Liberty Blue Automated Microwave Peptide Synthesizer with the manufacturers standard coupling cycles at 0.1 mmol scale. The purity of final compounds was confirmed by Waters LC-MS system. Systems were run with 0-40% methanol/water gradient with 0.1% TFA. All the purity of target compounds showed >95%.

*α-bromo peptides (G*PKRIA)*. Active bromo acetic acid (139mg, 1mmol) by DIC (155*μ*L, 1mmol) for 20min. Then added the solution into the peptide PKRIA on resin (0.1 mmol). The mixture was shaken overnight at r.t. The suspension was filtered, and the resin was washed with DMF (3ml x 3). Repeated the procedures again. Do test cleavage (TFA: TIPS: H_2_O=95:2.5:2.5) and MALDI-MS to check the progress of the reaction.

NAH-C2-GPKRIA (**1**). To a suspension of G*PKRIA on resin (0.1 mmol) in DMF (3 mL) was added compound **8** (91mg, 0.15 mmol) and DIPEA (26 *μ*L, 0.15 mmol). The mixture was shaken at room temperature overnight. The suspension was filtered, and the resin was subsequently washed with DMF (3 mL x 3), MeOH (3 mL x 3), and CH_2_Cl_2_ (3 mL × 3). The peptide conjugates were cleaved from the resin in the presence of a cleavage cocktail (10 mL) containing trifluoroacetic acid (TFA)/2,2’-(Ethylenedioxy)-diethanethiol (DODT) /triisopropylsilane (TIPS) /water (94:2.5:1:2.5 v/v) at room temperature for 4-5 h and resin was rinsed with a small amount of TFA. Volatiles of the filtrate were removed under N2 and the residue was precipitated with 10 vol. cold anhydrous ether, and collected by centrifugation. The supernatant was discarded and the pellet was washed with chilled ether and air-dried. The white precipitate was dissolved in ddH_2_O and purified by reverse phase HPLC using a waters system with a XBridge (BEH C18, 5μm, 10 × 250 mm) column with 0.1% TFA in water (A) and MeOH (B) as the mobile phase with monitoring at 214 nm and 254 nm. MALDI-MS (positive) m/z: calcd for C_44_H_77_N_18_O_11_ [M + H]^+^ m/z 1033.6019, found m/z 1033.7422.

NAH-C3-GPKRIA (**2**). Compound was prepared according to the procedure for NAH-C2-GPKRIA (**1**) and purified by reverse phase HPLC. MALDI-MS (positive) m/z: calcd for C_45_H_79_N_18_O_11_ [M + H]^+^ m/z 1047.6176, found m/z 1047.8483. NAH-C4-GPKRIA (**3**). Compound was prepared according to the procedure for NAH-C2-GPKRIA (**1**) and purified by reverse phase HPLC. MALDI-MS (positive) m/z: calcd for C_46_H_81_N_18_O_11_ [M + H]^+^ m/z 1061.6332, found m/z 1062.0251. NAH-C5-GPKRIA (**4**). Compound was prepared according to the procedure for NAH-C2-GPKRIA (**1**) and purified by reverse phase HPLC. MALDI-MS (positive) m/z: calcd for C_46_H_81_N_18_O_11_ [M + H]^+^ m/z 1075.6489, found m/z 1075.6666.

Tert-butyl (S)-4-((((3aR,4R,6S,6aR)-6-(6-amino-9H-purin-9-yl)-2,2-dimethyltetrahydrofuro[3,4-d][1,3]dioxol-4-yl)methyl)(4-(((benzyloxy)carbonyl)amino)butyl)amino)-2-((tert-butoxycarbonyl)amino)butanoate (**7a**). To a solution of tert-butyl (S)-4-((((3aR,4R,6S,6aR)-6-(6-amino-9H-purin-9-yl)-2,2-dimethyltetrahydrofuro[3,4-d][1,31dioxol-4-yl)methyl)amino)-2-((tert-butoxycarbonyl)amino)butanoate (**6**, 0.5g, 0.89 mmol) in methanol, benzyl (4-oxobutyl)carbamate (**5a**, 0.26g, 1.15 mmol) was add into the reaction. The mixture stirred for 20 min at r.t.. Then, NaBH_3_CN (82mg, 1.34 mmol) was added in protions and the reaction continued to stir for 6h at r.t.. After removing the solvent in vacuum, the residue was purified by column chromatography (CH_2_Cl_2_: MeOH = 30:1) to obtain 0.46g (68%) white solid. ^1^H NMR (500 MHz, Methanol-*d*_4_) δ 8.25 (s, 1H), 8.22 (s, 1H), 7.39 – 7.24 (m, 5H), 6.16 (d, *J* = 2.4 Hz, 1H), 5.53 – 5.49 (m, 1H), 5.05 (s, 2H), 5.04 – 5.01 (m, 1H), 4.37 – 4.28 (m, 1H), 4.09 – 4.02 (m, 1H), 3.14 – 3.00 (m, 2H), 2.89 – 2.75 (m, 1H), 2.68 – 2.54 (m, 2H), 2.53 – 2.31 (m, 3H), 1.95 – 1.84 (m, 1H), 1.72 – 1.61 (m, 1H), 1.58 (s, 3H), 1.45 – 1.44 (m, 2H), 1.43 (s, 9H), 1.42 (s, 9H), 1.38 (s, 3H).

Tert-butyl (S)-4-((5-((((9H-fluoren-9-yl)methoxy)carbonyl)amino)pentyl)(((3aR,4R,6S,6aR)-6-(6-amino-9H-purin-9-yl)-2,2-dimethyltetrahydrofuro[3,4-d][1,31dioxol-4-yl)methyl)amino)-2-((tert-butoxycarbonyl)amino)butanoate (**7b**). To a solution of tert-butyl (S)-4-((((3aR,4R,6S,6aR)-6-(6-amino-9H-purin-9-yl)-2,2-dimethyltetrahydrofuro[3,4-d][1,31dioxol-4-yl)methyl)amino)-2-((tert-butoxycarbonyl)amino)butanoate (**6**, 0.17g, 0.30 mmol) in methanol, (9H-fluoren-9-yl)methyl (5-oxopentyl)carbamate (**5b**, 0.13g, 0.40 mmol) was add into the reaction. The mixture stirred for 20 min at r.t.. Then, NaBH_3_CN (26mg, 0.42 mmol) was added in portion and the reaction continued to stir overnight at r.t.. After removing the solvent in vacuum, the residue was purified by column chromatography (CH_2_Cl_2_: MeOH = 50:1) to obtain 0.1g (38%) white solid. ^1^H NMR (500 MHz, Chloroform-*d*) δ 8.34 (s, 1H), 7.93 (s, 1H), 7.76 (d, *J* = 7.5 Hz, 2H), 7.60 (d, *J* = 6.9 Hz, 2H), 7.39 (t, *J* = 7.4 Hz, 2H), 7.30 (t, *J* = 7.3 Hz, 2H), 6.09 – 6.01 (m, 1H), 5.67 – 5.43 (m, 3H), 5.03 – 4.91 (m, 1H), 4.49 – 4.32 (m, 3H), 4.31 – 4.08 (m, 2H), 3.31 – 3.01 (m, 2H), 2.83 – 2.21 (m, 6H), 1.99 – 1.84 (m, 1H), 1.70 (ws, 3H), 1.61 (s, 3H), 1.44 (s, 18H), 1.39 (s, 3H), 1.34 – 1.10 (m, 4H).

Tert-butyl (S)-4-((((3aR,4R,6S,6aR)-6-(6-amino-9H-purin-9-yl)-2,2-dimethyltetrahydrofuro[3,4-d][1,3]dioxol-4-yl)methyl)(4-aminobutyl)amino)-2-((tert-butoxycarbonyl)amino)butanoate (**10**). To a solution of **7a** (0.46g, 0.6mmol) in MeOH, 10 % Pd/C was added slowly into the reaction. The mixture was degassed under vacuum and then under H2 atmosphere and stirred overnight. After filtration and concentration, the residue was purified by column chromatography (CH_2_Cl_2_: MeOH = 10:1) to get 0.21g (56%) white solid. ^1^H NMR (500 MHz, Methanol-*d*_4_) δ 8.26 (s, 1H), 8.22 (s, 1H), 6.17 (d, *J* = 2.1 Hz, 1H), 5.54 – 5.51 (m, 1H), 5.08 – 5.01 (m, 1H), 4.37 – 4.26 (m, 1H), 4.13 – 4.00 (m, 1H), 2.84 – 2.74 (m, 1H), 2.66 – 2.52 (m, 4H), 2.50 – 2.39 (m, 2H), 2.38 – 2.28 (m, 1H), 2.00 – 1.86 (m, 1H), 1.69 – 1.60 (m, 1H), 1.59 (s, 3H), 1.46 – 1.41 (m, 20H), 1.38 (s, 3H).

Tert-butyl (S)-4-((((3aR,4R,6S,6aR)-6-(6-amino-9H-purin-9-yl)-2,2-dimethyltetrahydrofuro[3,4-d][1,3]dioxol-4-yl)methyl)(5-aminopentyl)amino)-2-((tert-butoxycarbonyl)amino)butanoate (**11**). To a solution of **7b** (0.1g, 0.11mmol) in CH_3_CN cooled in ice-bath, DBU (34 *μ*L, 0.22 mmol) was added in dropwise. The reaction stirred for 1h at r.t.. The solvent was removed in vacuum, and the residue was purified by column chromatography (CH_2_Cl_2_: MeOH = 10:1) to get 50 mg (71%) product. ^1^H NMR (500 MHz, Methanol-*d*_4_) δ 8.26 (s, 1H), 8.22 (s, 1H), 6.17 (d, *J* = 2.2 Hz, 1H), 5.57 – 5.50 (m, 1H), 5.08 – 5.02 (m, 1H), 4.36 – 4.28 (m, 1H), 4.11 – 4.01 (m, 1H), 2.84 – 2.74 (m, 1H), 2.66 – 2.54 (m, 4H), 2.48 – 2.39 (m, 2H), 2.38 – 2.29 (m, 1H), 1.99 – 1.88 (m, 1H), 1.68 – 1.61 (m, 1H), 1.59 (s, 3H), 1.44 (s, 9H), 1.44 (s, 9H), 1.42 – 1.40 (m, 2H), 1.39 (s, 3H), 1.37 – 1.30 (m, 2H), 1.30 – 1.17 (m, 2H).

### Protein Expression and Purification

Expression and purification of NTMT1, G9a, SETD7, PRMT1, PRMT3, *Tb*PRMT7 and NNMT was performed as previously described ^[16],[18],[35],[37]–[40]^

### NTMT1 Biochemical Assays and Enzyme Kinetics Study

A fluorescence-based SAHH-coupled assay was applied to study the IC_50_ values of compounds with both SAM and the peptide substrate at their *K*_m_ values. For 10min pre-incubation, the methylation assay was performed under the following conditions in a final well volume of 40 *μ*L: 25 mM Tris (pH = 7.5), 50 mM KCl, 0.01% Triton X-100, 5 *μ*M SAHH, 0.1 *μ*M NTMT1, 3 *μ*M SAM, and 10 *μ*M ThioGlo4. The inhibitors were added at concentrations ranging from 0.15 nM to 10 *μ*M. After 10 min incubation, reactions were initiated by the addition of 0.5 *μ*M GPKRIA peptide. For 120min pre-incubation, NTMT1 and compounds were pre-incubated for 120min. Then the pre-incubation mixture was added into reaction mixture: 25 mM Tris (pH = 7.5), 50 mM KCl, 0.01% Triton X-100, 5 *μ*M SAHH, 3 *μ*M SAM, and 10 *μ*M ThioGlo4. The reactions were initiated by the addition of 0.5 *μ*M GPKRIA peptide. Fluorescence was monitored on a BMG CLARIOstar microplate reader with excitation 400 nm and emission 465 nm. Data were processed by using GraphPad Prism software 7.0.

For the inhibition mechanism studies, varying concentrations of SAM with 0.5 *μ*M fixed concentration of GPKRIA or varying concentration of GPKRIA with 3 *μ*M fixed concentration of SAM were included in reactions at concentration of inhibitors ranging from 0.15 nM to 10 *μ*M. Fluorescence was monitored on a BMG CLARIOstar microplate reader with excitation 400 nm and emission 465 nm. All experiments were determined in duplicate. Data were processed by using GraphPad Prism software 7.0.

### ITC

Isothermal titration calorimetry (ITC) measurements were performed by VP-ITC MicroCal calorimeter at 25 °C. Titration buffer contains 20 mM Tris-HCl pH 7.5, 150 mM NaCl and 0.5 mM TECP (tris(2-carboxyethyl)phosphine). Purified NTMT1 and NTMT2 were diluted with the ITC buffer to the final concentration of 30-50 *μ*M, and the compounds were dissolved to 0.5-1mM in the same buffer. The compound was titrated into the protein solution with 26 injections of 10 μl each. Injections were spaced 180 sec with a reference power of 15 μcal/sec. The ITC data were processed with Origin 7.0 software (Microcal) using one-site binding or two-site binding model.

### Time-dependent Inhibition Study

The fluorescence-based SAHH-coupled assay was applied to this study. The assay was performed under the following conditions in a final well volume of 40 *μ*L: 25 mM Tris (pH = 7.5), 50 mM KCl, 0.01% Triton X-100, 5 *μ*M SAHH, 0.1 *μ*M NTMT1, 3 *μ*M SAM, 0.5 *μ*M GPKRIA and 10 *μ*M ThioGlo4. The inhibitors were added at concentrations ranging from 0.15 nM to 10 *μ*M. NTMT1 and compounds were pre-incubated for 30min, 60min, 90min, and 120min, Then the preincubation mixture was added into reaction mixture: 25 mM Tris (pH = 7.5), 50 mM KCl, 0.01% Triton X-100, 5 *μ*M SAHH, 3 *μ*M SAM, and 10 *μ*M ThioGlo4. The reactions were initiated by the addition of 0.5 *μ*M GPKRIA peptide. Fluorescence was monitored on a BMG CLARIOstar microplate reader with excitation 400 nm and emission 465 nm. Data were processed by using GraphPad Prism software 7.0 with Morrison’s quadratic equation.

### Selectivity Assays

A fluorescence-based SAHH-coupled assay was applied to study the effect of compounds on methyltransferase activity of G9a, SETD7, PRMT1, PRMT3 and *Tb*PRMT7. For G9a, the assay was performed in a final well volume of 40 *μ*L: 25 mM potassium phosphate buffer (pH = 7.6), 1 mM EDTA, 2 mM MgCl_2_, 0.01% Triton X-100, 5 *μ*M SAHH, 0.1 *μ*M His-G9a, 10 *μ*M SAM, and 10 *μ*M ThioGlo4. The inhibitors with varied concentration were added. After 10 min incubation with inhibitor**s**, reactions were initiated by the addition of 4 *μ*M H3-21 peptide. Fluorescence was monitored on a BMG CLARIOstar microplate reader with excitation 400 nm and emission 465 nm. Data were processed by using GraphPad Prism software 7.0.

For SETD7, the assay was performed in a final well volume of 40 *μ*L: 25 mM potassium phosphate buffer (pH = 7.6), 0.01% Triton X-100, 5 *μ*M SAHH, 1 *μ*M His-SETD7, 2 *μ*M SAM, and 15 *μ*M ThioGlo1. After 10 min incubation with inhibitors, reactions were initiated by the addition of 90 *μ*M H3-21 peptide. For PRMT1, the assay was also performed in a final well volume of 40 *μ*L: 2.5 mM HEPES (pH = 7.0), 25 mM NaCl, 25 *μ*M EDTA, 50 *μ*M TCEP, 0.01% Triton X-100, 5 *μ*M SAHH, 0.2 *μ*M PRMT1, 10 *μ*M SAM, and 15 *μ*M ThioGlo1. After 10 min incubation with inhibitors, reactions were initiated by the addition of 5 *μ*M H4-21 peptide. For PRMT3, the assay was also performed in a final well volume of 40 *μ*L: 2.5 mM HEPES (pH = 7.0), 25 mM NaCl, 25 *μ*M EDTA, 50 *μ*M TCEP, 0.01% Triton X-100, 5 *μ*M SAHH, 1 *μ*M PRMT3, 25 *μ*M SAM, and 15 *μ*M ThioGlo1. After 10 min incubation with inhibitors, reactions were initiated by the addition of 10 *μ*M H4-21 peptide. For *Tb*PRMT7, the assay was performed in a final well volume of 40 *μ*L: 25 mM Tris (pH = 7.5), 50 mM KCl, 0.01% Triton X-100, 5 *μ*M SAHH, 0.2 *μ*M

*Tb*PRMT7, 3 *μ*M SAM, and 15 *μ*M ThioGlo1. After 10 min incubation with inhibitors, reactions were initiated by the addition of 60 *μ*M H4-21 peptide. For SAHH, the assay was also performed in a final well volume of 40 *μ*L: 25 mM Tris (pH = 7.5), 50 mM KCl, 0.01% Triton X-100, 0.1 *μ*M SAHH and 15 *μ*M ThioGlo1. After 10min incubation with compounds, 0.5 *μ*M SAH was added to initiate the reactions. All experiments were determined in duplicate. Fluorescence was monitored on a BMG CLARIOstar microplate reader with excitation 380 nm and emission 505 nm.

For NNMT, a noncoupled fluorescence assay was performed in a final well volume of 40 *μ*L as described before^[41],[42]^: 5 mM potassium phosphate buffer (pH = 7.6), 0.4 *μ*M NNMT and 20 *μ*M SAM. After 10min incubation with compounds, 25 *μ*M 1-quinoline was added to initiate the reactions. All experiments were determined in duplicate. Fluorescence was monitored on the BMG CLARIOstar microplate reader with excitation 330 nm and emission 405 nm.

### MALDI-MS Methylation Assay and Methylation Inhibition Assay

MALDI-MS methylation and inhibition assay were performed and analyzed via a Sciex 4800 MALDI TOF/TOF MS. MALDI-MS methylation assay was performed in a final well volume of 40 *μ*L: 0.1 *μ*M NTMT1, 20 mM Tris (pH = 7.5), 50 mM KCl, 3 *μ*M SAM and 10 *μ*M compounds. After incubation overnight at 37 °C, the samples were quenched in a 1:1 ratio with a quenching solution (20 mM NH_4_H_2_PO_4_, 0.4% (v/v) TFA in 1:1 acetonitrile/water). Samples were analyzed by MALDI-MS with 2,5-dihydroxybenzoic acid matrix solution.

The inhibition assay was performed in a final well volume of 40 *μ*L: 0.1 *μ*M NTMT1, 20 mM Tris (pH = 7.5), 50 mM KCl, 3 *μ*M SAM and various concentrations of compounds at 37 °C for 10 min before the addition of 0.5 *μ*M GPKRIA peptide to initiate the reaction. After incubation for 20 min, the samples were quenched in a 1:1 ratio with a quenching solution (20 mM NH_4_H_2_PO_4_, 0.4% (v/v) TFA in 1:1 acetonitrile/water). Samples were analyzed by MALDI-MS with 2,5-dihydroxybenzoic acid matrix solution. Duplicate was performed for all experiments. Data were processed in Data Explorer.

### Co-crystallization and Structure Determination

For co-crystallization, NTMT1 was mixed with 2 mM compounds incubated for 1 hr at 4°C, diluted 6-fold in 1x TBS, pH 7.6, with 1 mM TCEP, and then re-concentrated to 30 mg/mL. Broad matrix and optimization crystallization screening were performed using a Mosquito-LCP high throughput crystallization robot (TTP LabTech) using hanging-drop vapor diffusion method, incubated at 20°C, and crystals harvested directly from the 96-well crystallization plates. Crystals containing **1** were grown in 32 % (w/v) PEG 4000, 0.8 M Lithium Chloride, 0.1 M TrisoHCl (pH = 8.5), and crystals containing **4** were grown in 30% (w/v) PEG3350, 0.2 M Li_2_SO_4_, 0.1 M bis-tris propane (pH = 7.0); all crystals were harvested directly from the drop and flash cooled into liquid nitrogen.

Data were collected on single crystals at 12.0 keV at the GM/CA-CAT ID-D beamline at the Advanced Photon Source, Argonne National Laboratory. The data were processed using the on-site automated pipeline with Fast DP and molecular replacement performed with Phaser-MR (PHENIX)^[43],[44]^. All model building was performed using COOT and subsequent refinement done using phenix.refine (PHENIX)^[44],[45]^. Structure factors and model coordinates have been deposited in the Protein Data Bank with PDB IDs: 6WJ7 (**1**) and 6PVB (**4**). Structure-related figures were made with PyMOL (Schrödinger) and annotated and finalized with Adobe Photoshop and Illustrator.

## Acknowledgements

We appreciate Dr. Darrel L. Peterson for purification of SAHH. We thank Krystal Diaz for purifying *Tb*PRMT7. The authors acknowledge the support from NIH grants R01GM117275 (RH), 1R01GM127896 (NN), 1R01AI127793 (NN), and P30 CA023168 (Purdue University Center for Cancer Research). The SGC is a registered charity (number 1097737) that receives funds from AbbVie, Bayer Pharma AG, Boehringer Ingelheim, Canada Foundation for Innovation, Eshelman Institute for Innovation, Genome Canada through Ontario Genomics Institute [OGI-055], Innovative Medicines Initiative (EU/EFPIA) [ULTRA-DD grant no. 115766], Janssen, Merck KGaA, Darmstadt, Germany, MSD, Novartis Pharma AG, Ontario Ministry of Research, Innovation and Science (MRIS), Pfizer, São Paulo Research Foundation-FAPESP, Takeda, and Wellcome. We also thank supports from the Department of Medicinal Chemistry and Molecular Pharmacology (RH) and Department of Biological Sciences (NN) at Purdue University.

## Table of Contents

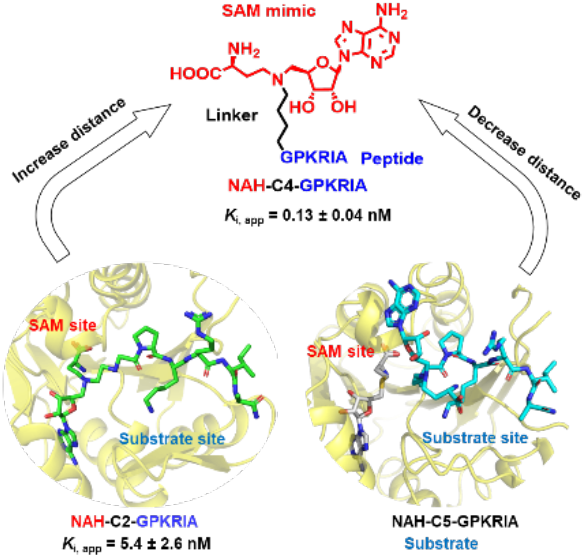

A series of bisubstrate analogs were examined to probe the protein N-terminal methyltransferase 1 (NTMT1) active site through biochemical characterization and structural studies. A 4-C atom linker yields the most potent bisubstrate inhibitor for NTMT1, while a 5-C atom linked bisubstrate analog functions as a substrate instead of an inhibitor.

