## supplementary for "Probing the Plasticity in the Active Site of Protein N-terminal Methyltransferase 1 Using Bisubstrate Analogs"

### Table of Contents

A. NMR and MS spectra of compound **7a-b**, **10-11**

B. NMR, MALDI-MS and LC-MS/HPLC spectra of compound **1-4**

C. Table S1. $IC_{50}$ values of bisubstrate compounds.

D. Table S2. Selectivity evaluation of compound **3**

E. Table S3. Crystallography data and refinement statistics (PDB ID: 6WJ7 and 6PVB).

**NMR spectra of compound 7a**

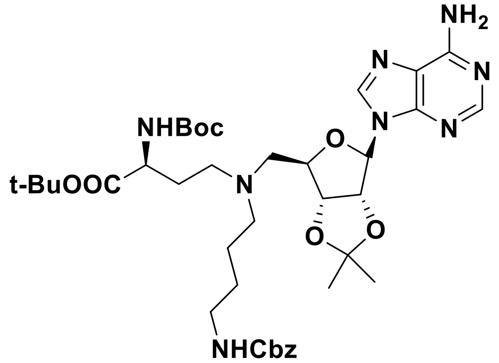

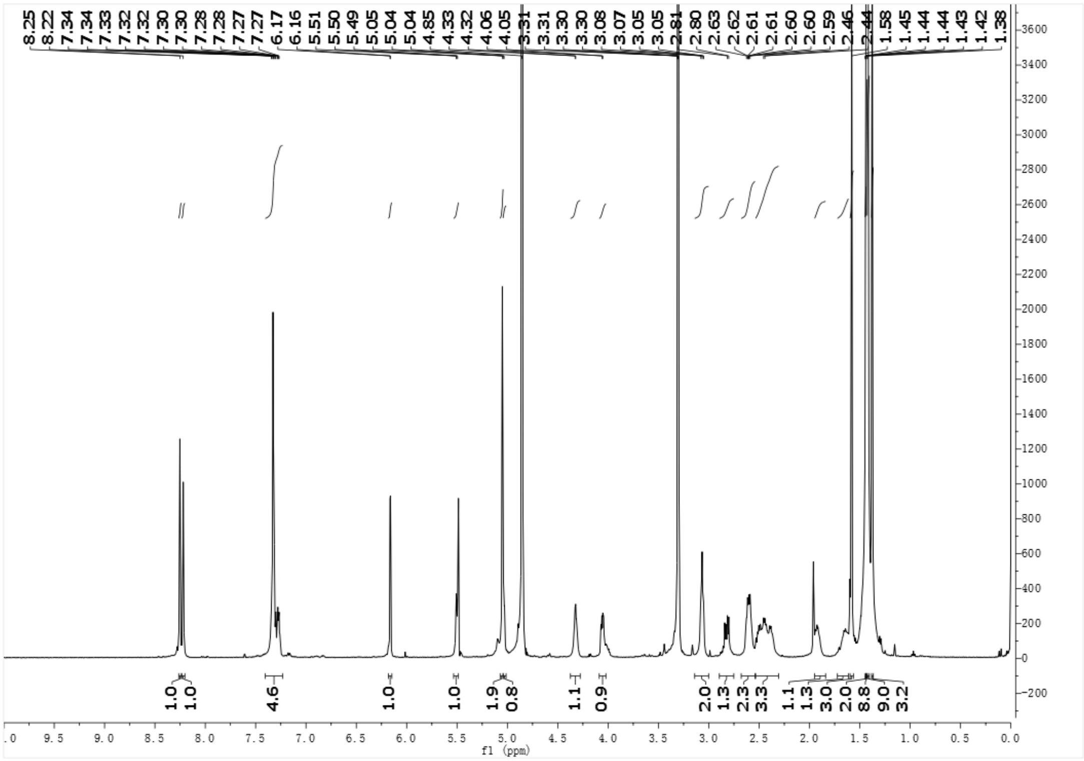

**NMR spectra of compound 7b**

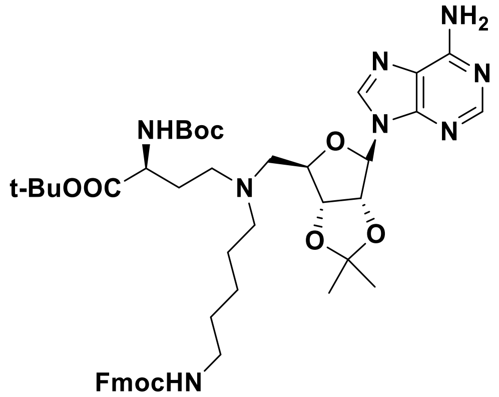

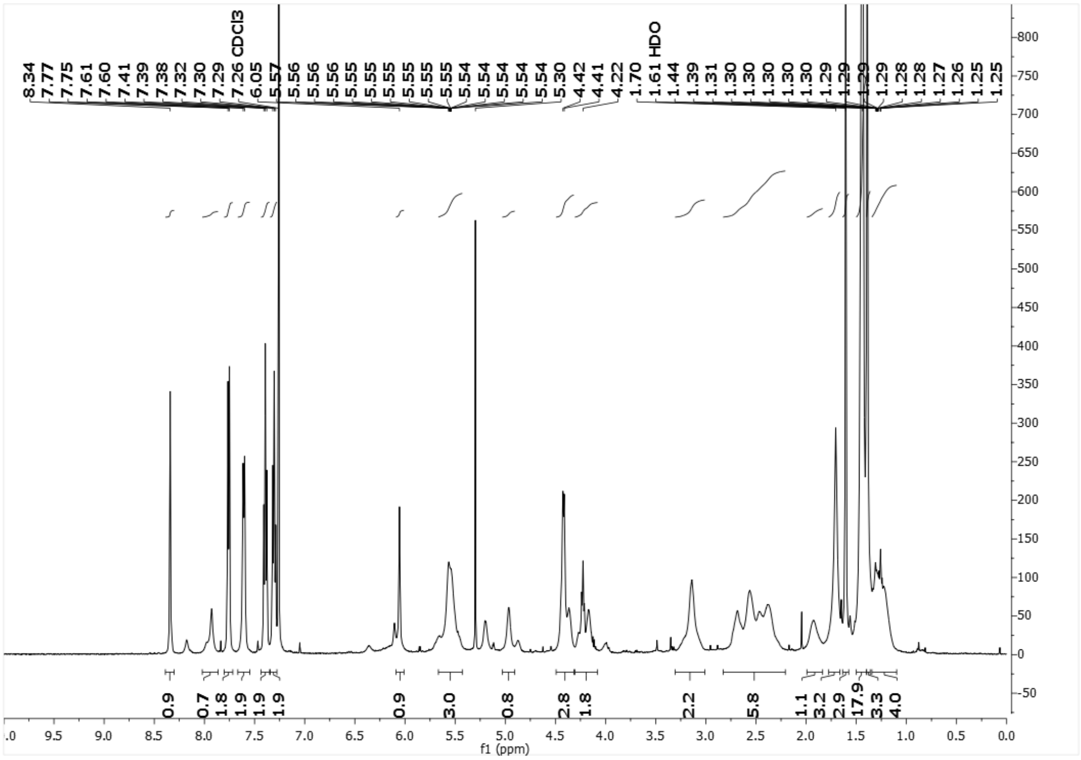

**NMR spectra of compound 10**

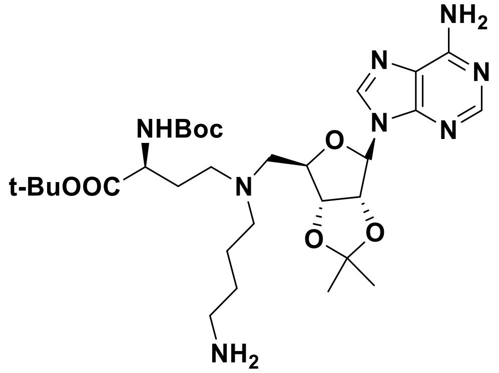

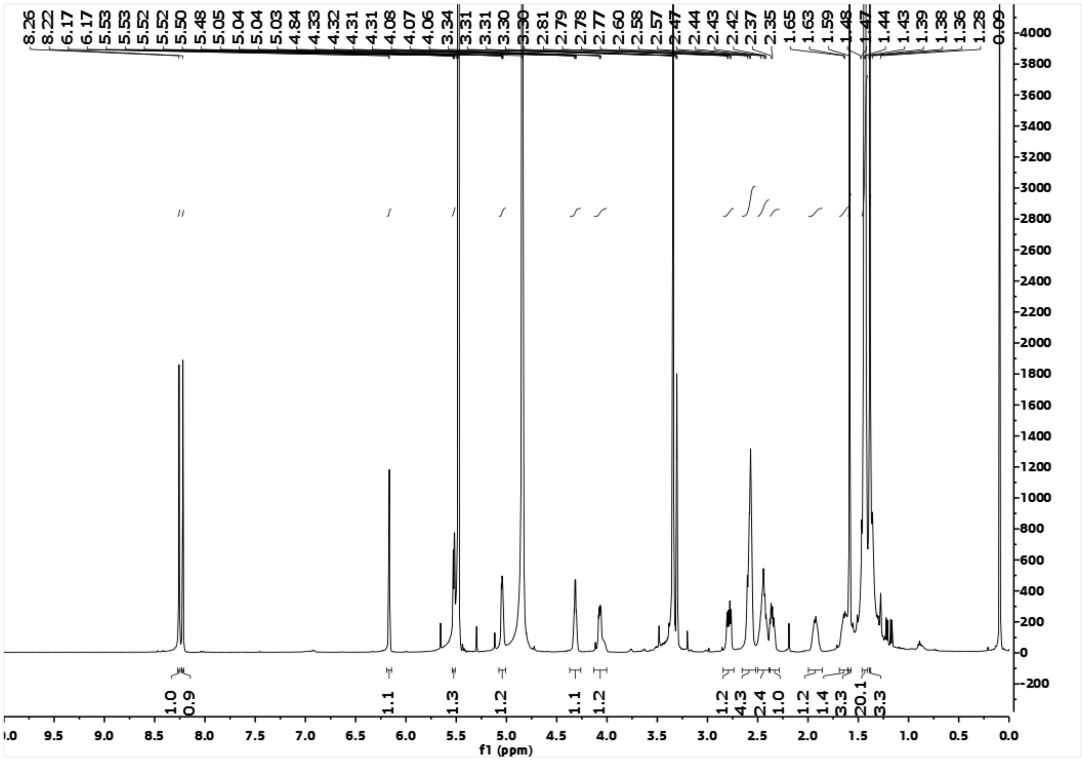

**NMR and MS spectra of compound 11**

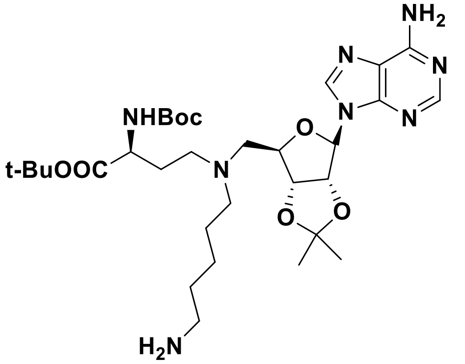

NMR spectra of compound **11**

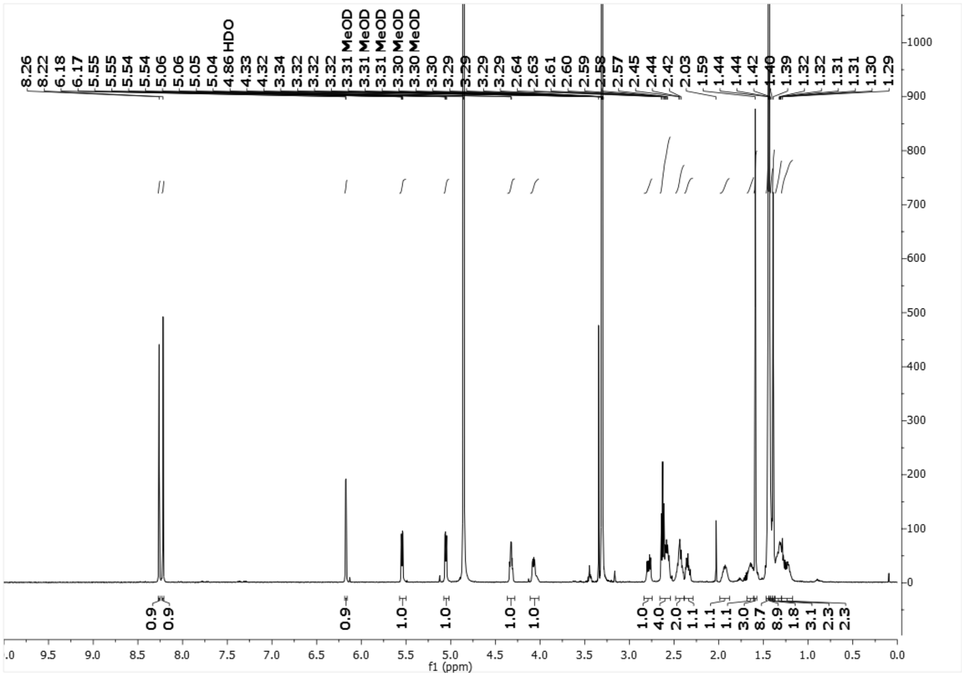

MS spectra of compound **11**

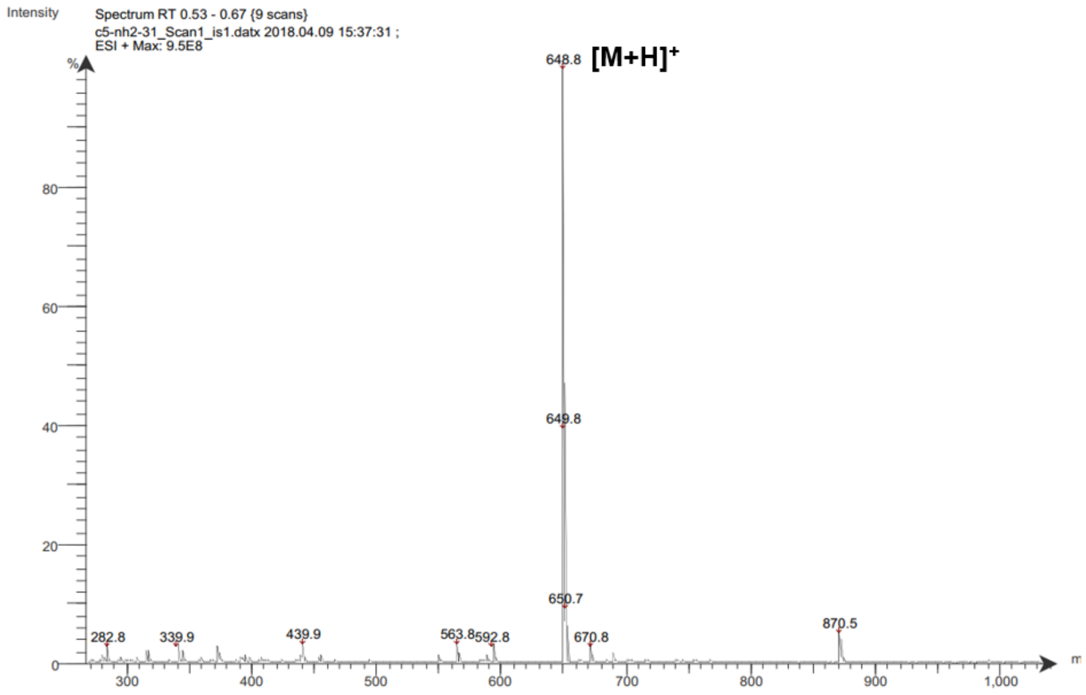

**MALDI-MS and LC-MS spectra of compound 1**

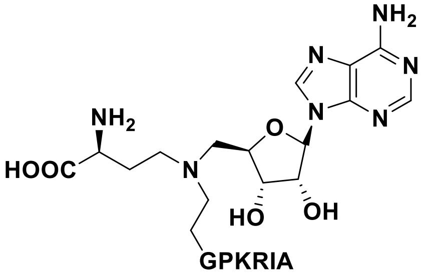

**NAH-C2-GPKRIA**

MALDI-MS spectra of compound **1**

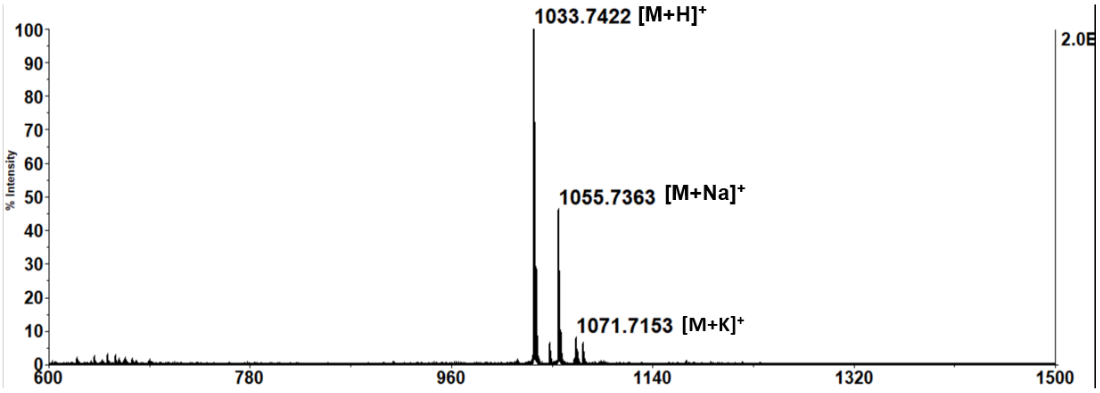

LC-MS spectra of compound **1**

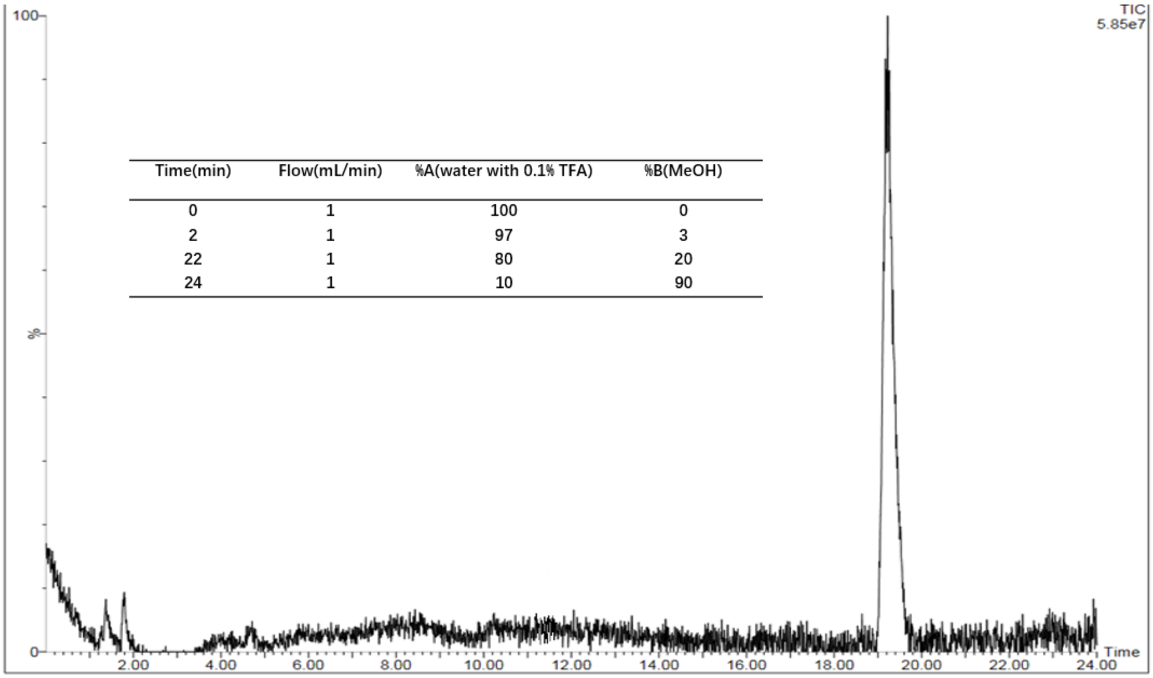

| Time(min) | Flow(mL/min) | %A(water with 0.1% TFA) | %B(MeOH) |
|---|---|---|---|
| 0 | 1 | 100 | 0 |
| 2 | 1 | 97 | 3 |
| 22 | 1 | 80 | 20 |
| 24 | 1 | 10 | 90 |

**MALDI-MS and LC-MS spectra of compound 2**

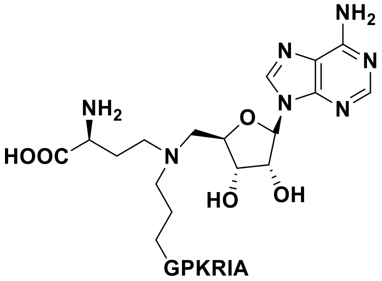

NAH-C3-GPKRIA

MALDI-MS spectra of compound **2**

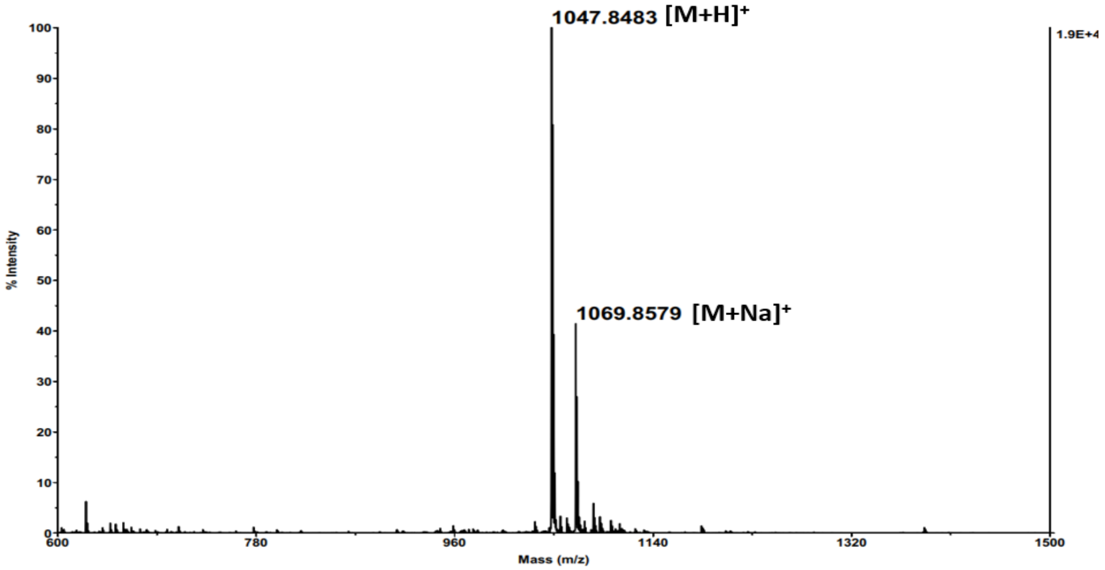

LC-MS spectra of compound **2**

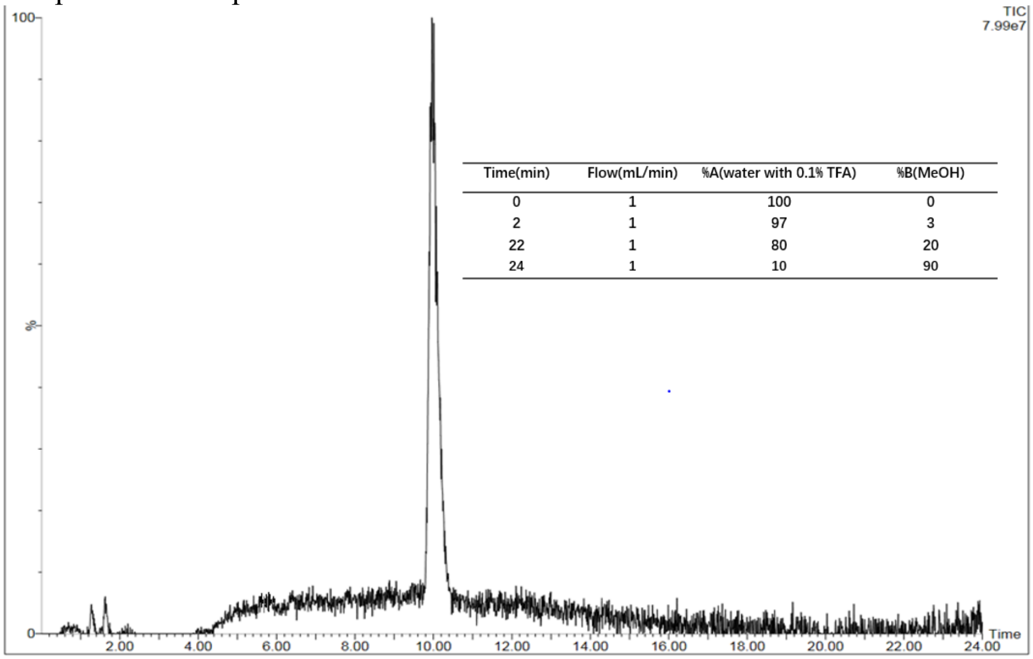

| Time(min) | Flow(mL/min) | %A(water with 0.1% TFA) | %B(MeOH) |
|---|---|---|---|
| 0 | 1 | 100 | 0 |
| 2 | 1 | 97 | 3 |
| 22 | 1 | 80 | 20 |
| 24 | 1 | 10 | 90 |

**MALDI-MS and LC-MS spectra of compound 3**

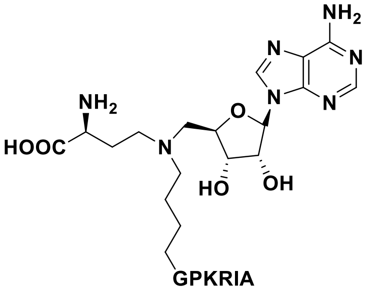

NAH-C4-GPKRIA

MALDI-MS spectra of compound **3**

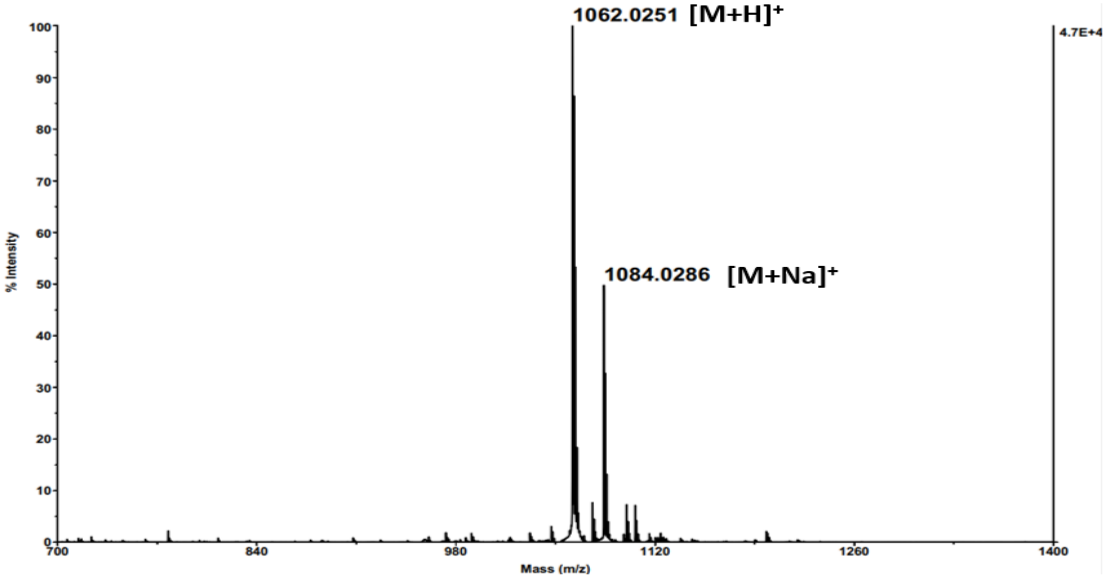

LC-MS spectra of compound **3**

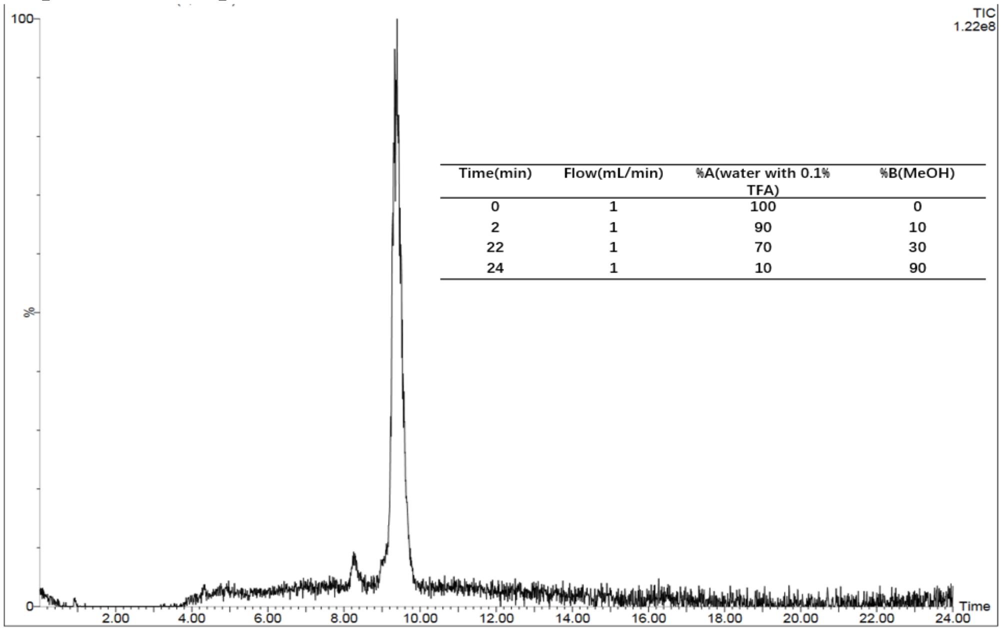

| Time(min) | Flow(mL/min) | %A(water with 0.1% TFA) | %B(MeOH) |
|---|---|---|---|
| 0 | 1 | 100 | 0 |
| 2 | 1 | 90 | 10 |
| 22 | 1 | 70 | 30 |
| 24 | 1 | 10 | 90 |

**MALDI-MS and LC-MS spectra of compound 4**

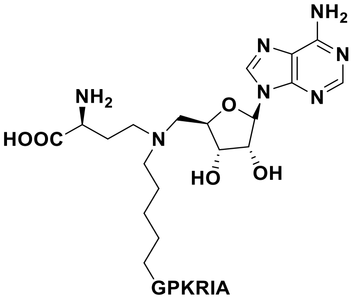

NAH-C5-GPKRIA

MALDI-MS spectra of compound **4**

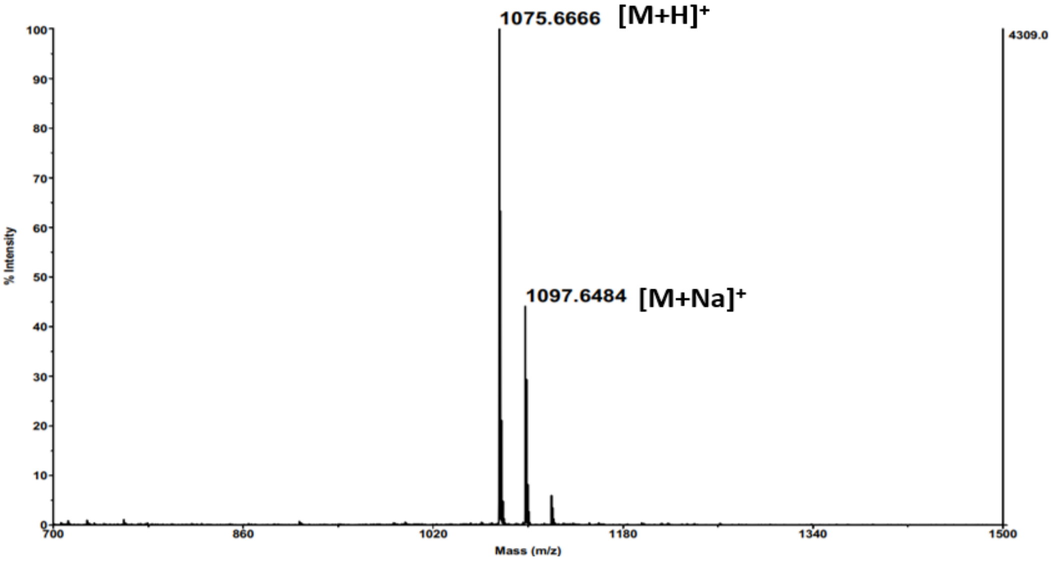

LC-MS spectra of compound **4**

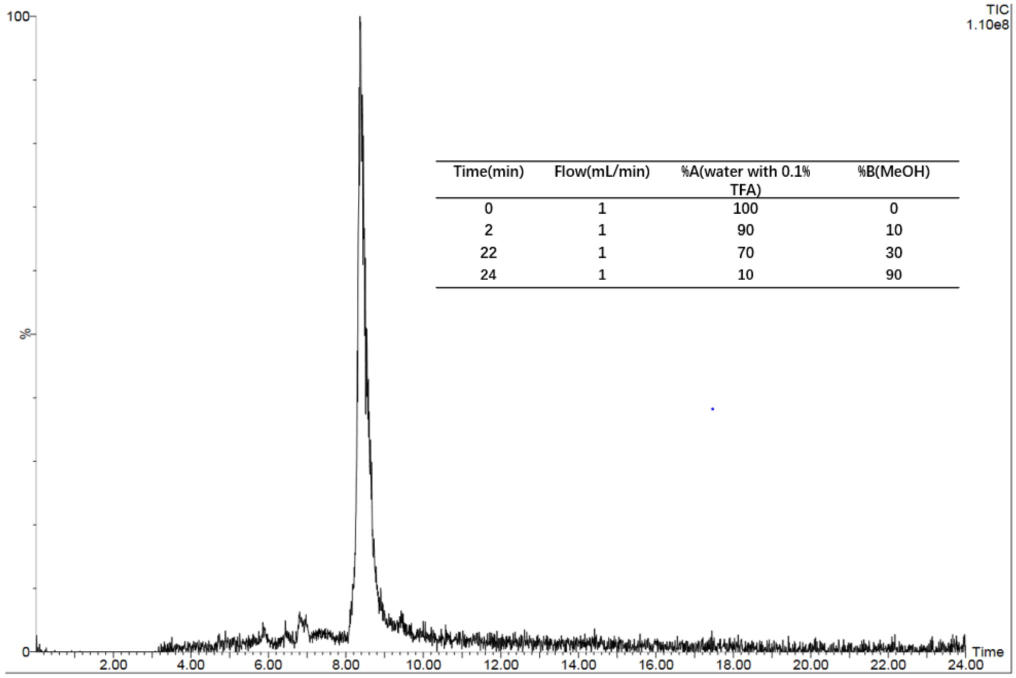

| Time(min) | Flow(mL/min) | %A(water with 0.1% TFA) | %B(MeOH) |
|---|---|---|---|
| 0 | 1 | 100 | 0 |
| 2 | 1 | 90 | 10 |
| 22 | 1 | 70 | 30 |
| 24 | 1 | 10 | 90 |

**Table S1. $IC_{50}$ values of bisubstrate compounds**

| Compound ID | $IC_{50}$ (nM)[a] |
| --- | --- |
| **1 (NAH-C2-GPKRIA)** | 260±70 |
| **2 (NAH-C3-GPKRIA)** | 130±40 |
| **3 (NAH-C4-GPKRIA)** | 82±17 |
| **4 (NAH-C5-GPKRIA)** | N.A. |

[a] The values of $IC_{50}$ for all compounds were determined with triplicate experiments (n = 3). N.A.: not applied.

**Table S2. Selectivity evaluation of compound 3**

| Enzyme activity (%) [a] | [**3**] ($\mu$M) | | | | $IC_{50}$($\mu$M) |
|---|---|---|---|---|---|
| | 3.7 | 11 | 33 | 100 | |
| G9a | 95 | 94 | 88 | 78 | >100 |
| SETD7 | 101 | 106 | 101 | 86 | >100 |
| PRMT1 | 98 | 93 | 78 | 47 | >33 |
| PRMT3 | 98 | 87 | 58 | 26 | >33 |
| *Tb*PRMT7 | 93 | 82 | 60 | 30 | >33 |
| NNMT | 99 | 97 | 97 | 93 | >100 |
| SAHH | 84 | 71 | 57 | 40 | >33 |

[a] The values of enzyme activity are mean values of duplicate experiments (n = 2).

**Table S3. Crystallography data and refinement statistics.**

| **Data Collection** | **NTMT1-NAH-C2-GPKRIA** | **NTMT1-NAH-C5-GPKRIA** |
|---|---|---|
| λ (Å) | 0.97934 | 1.0332 |
| Space group | $P3_121$ | $P3_121$ |
| a, b, c (Å) | 72.80, 72.80, 80.13 | 72.95, 72.95, 82.09 |
| α, β, γ (°) | 90, 90, 120 | 90, 90, 120 |
| Resolution (Å)* | 50 - 1.42 (1.44 - 1.42) | 30 - 1.50 (1.53 - 1.50) |
| Completeness (%)* | 99.93 (99.78) | 99.8 (98.3) |
| Redundancy* | 6.5 (9.3) | 5.3 (5.3) |
| $R_{sym}$ †* | 0.12 (1.3) | 0.05 (3.3) |
| I / σ (I)* | 20.4 (1.0) | 14.5 (0.4) |
| $CC_{1/2}$ | 0.99 (0.46) | 1.0 (0.17) |
| **Refinement** | | |
| Resolution (Å) | 24.59 – 1.42 | 30 – 1.50 |
| No. reflections | 46403 | 41,064 |
| R§/$R_{free}$¶ | 0.16/0.18 | 0.21/0.23 |
| **r.m.s. deviations** | | |
| Bonds (Å) | 0.015 | 0.003 |
| Angles (°) | 1.702 | 0.827 |
| No. Protein atoms | 1743 | 1858 |
| No. Ligand atoms | 73 | 102 |
| No. Waters | 233 | 182 |
| **B-factors** ($Å^2$) | | |
| Wilson B | 16.51 | 28.86 |
| Protein | 23.31 | 37.09 |
| Ligands | 26.50 | 50.88 |
| Waters | 37.19 | 45.22 |
| **Ramachandran Analysis**¥ | | |
| Favored (%) | 96.77 | 98.2 |
| Allowed (%) | 2.76 | 1.8 |
| Outliers (%) | 0.46 | 0 |
| PDB code | 6WJ7 | 6PVB |

† $R_{sym} = \Sigma_{hkl,j}\,(|I_{hkl} - \langle I_{hkl}\rangle|) / \Sigma_{hkl,j}\, I_{hkl}$, where $\langle I_{hkl}\rangle$ is the average intensity for a set of j symmetry related reflections and $I_{hkl}$ is the value of the intensity for a single reflection within a set of symmetry-related reflections.

§ $R$ factor $= \Sigma_{hkl}\,(||F_o| - |F_c||) / \Sigma_{hkl}|F_o|$ where $F_o$ is the observed structure factor amplitude and $F_c$ is the calculated structure factor amplitude.

¶ $R_{free} = \Sigma_{hkl,T}\,(||F_o| - |F_c||) / \Sigma_{hkl,T}|F_o|$, where a test set, T (5% of the data), is omitted from the refinement.

¥ Performed using Molprobity within PHENIX.
* Indicates statistics for last resolution shell shown in parenthesis.
